# Corazonin links social role and metabolic-stress responses in honey bee workers

**DOI:** 10.64898/2026.10.06.757184

**Authors:** Sebastian L. Scofield, Gro V. Amdam

## Abstract

Corazonin is implicated in regulating diverse behavioral and physiological responses in insects, yet its role in honey bees remains unclear. Work in social and non-social species suggests an ancestral function for corazonin as a metabolic stress signal and a derived function as a pro-foraging signal in social insects. To investigate this hypothesis, we examined corazonin signaling across worker roles and metabolic states and tested its regulatory relationships with pathways central to worker division of labor. Central corazonin levels (both transcript and peptide) were approximately threefold higher in foragers than in nurses. Acute metabolic challenge did not alter brain corazonin but strongly upregulated fat body corazonin receptor (CrzR) transcript in nurses only, suggesting role-specific induction of peripheral sensitivity to corazonin signaling. Injection of synthetic corazonin decreased fat body insulin receptor substrate transcript, while transcripts of vitellogenin (Vg), unpaired-like A, ecdysone receptor B, and Cyp314a1 were unaffected. Measures of gustatory responsiveness, starvation resistance, and hemolymph sugar levels were similarly unaffected by corazonin injection. Reducing Vg through RNA interference in nurse-age bees did not affect CrzR transcript levels, despite vitellogenin’s established roles in stress buffering, antioxidant defense, and lifespan. This contrast highlights the specificity of CrzR upregulation in nurses as a response to metabolic challenge rather than general stress. Together, these findings support corazonin as a co-opted nutritional signal that helps shape the forager physiological state, suggesting that it may act in part through interactions with insulin signaling rather than directly regulating classical nutritional or stress-response pathways.

## Introduction

Neuropeptides are crucial regulators of many behavioral, developmental, and physiological functions throughout the life cycle of animals. This signaling can be very complex, acting over a wide range of temporal and spatial scales, and having pleiotropic functions: in some cases, the same neuropeptide can play roles both as a co-transmitter and/or neuromodulator in the brain, and circulating hormone in the periphery (Nässel and Zandawala 2019). This dual action can allow these neuropeptides to act as central players in brain-body communication, linking internal physiological state and behavioral output. One of the most critical needs for this communication is in response to metabolic stress, which requires coordination of behavior and physiology to maintain homeostasis and ensure the survival of the organism (Auer et al. 2020). Insects, with their less complex neuronal and endocrine systems relative to mammals, have emerged as useful model species for studying the role of neuropeptides in coordinating the organismal response to metabolic stress (Nässel and Zandawala 2019).

In insects, the brain-fat body axis is especially important for metabolic balance: the fat body is the central organ for energetic and nutritional sensing and storage, protein and lipid synthesis, and is the most metabolically active tissue in adults (Arrese and Soulages 2010). An effective stress response thus requires coordination of fat body metabolic response (such as mobilization of stored macronutrients) with a behavioral response (such as food-seeking). The neuropeptide corazonin is a hypothesized coordinator of the response to metabolic stress in insects (Veenstra 2009; Kubrak et al. 2016). Corazonin is found across many invertebrate taxa, where evidence shows it is involved in regulating diverse aspects of behavior, physiology, and development (Hamoudi et al. 2016; Arendt et al. 2017; Hou et al. 2017; Alexander et al. 2018; Imura et al. 2020; Khan et al. 2021; Xi et al. 2025; Zhang et al. 2025). Corazonin is perhaps best studied in the model organism *Drosophila*, where it is mainly synthesized in a set of neurosecretory cells in the brain and then released into the hemolymph for systemic circulation (Zandawala et al. 2021). It is also expressed in some non-neurosecretory neurons (McKim et al. 2024). The corazonin receptor (CrzR) is expressed in the brain, salivary glands, heart, and fat body of adult *Drosophila* (Kubrak et al. 2016), fitting with the function of corazonin as both central and peripheral regulator. During metabolic stress, corazonin signaling promotes locomotor activity and food intake; it also regulates fat body carbohydrate metabolism and expression of metabolic genes such as the adipokine Upd2 (Kubrak et al. 2016), which signals energetic state from fat body to brain (Rajan and Perrimon 2012). Fat body-specific knockdown of *CrzR* also produced downstream endocrine changes in the brain. This included altered expression of two peptide hormones in the insulin/insulin-like growth factor signaling (IIS) pathway, a central anabolic regulator of growth, development, and metabolism (Nässel et al. 2015), as well as substantially reduced adipokinetic hormone (AKH) brain immunoreactivity, suggesting increased release of AKH (Kubrak et al. 2016), a glucagon-like hormone that controls energy mobilization in insects and many other protostomes (Van der Horst 2003; Bednářová et al. 2013). In addition to these endocrine effects, recent connectomic work further shows that a subset of corazonin neurons directly innervate the wing neuropil, providing a pathway by which corazonin signaling can modulate flight-related motor activity (McKim et al. 2024). These links to known regulators of growth, metabolism, and behavior support the hypothesized function of corazonin as a signal of metabolic stress throughout insects.

Metabolic stress pathways are proposed to have evolved a novel role in eusocial insects as the fundamental organizing factor in colony division of labor (Negroni and LeBoeuf 2023; Walton and Toth 2023). This is very well studied in the queen-worker divide, where reproductive queens typically receive higher quality and more food than non-reproductive workers at every stage of development (Page Jr. and Amdam 2007). While less well recognized, these same metabolic stress pathways have been proposed as structuring the behavior and physiology of worker subcastes (Walton and Toth 2023). Workers have low or no reproductive output and so contribute to colony fitness through performing tasks that indirectly contribute to reproduction. In many eusocial insect species, tasks performed by workers are organized by division of workers into two main behavioral subcastes: nest workers engage in tasks such as caring for brood, feeding nestmates, and cleaning the nest, while foragers leave the relative safety of the nest and collect food to bring back to the colony (Toth and Robinson 2005; Silberman et al. 2016; Walton and Toth 2023). Optimal foraging theory predicts that natural selection will shape colony division of labor to maximize energy intake and minimize energy loss (Oster and Wilson 1978). Because foragers have higher mortality risk, colonies that allocate nutrient resources away from foragers towards castes with longer average lifespans should thus have higher fitness, and indeed a pattern of nest workers with higher nutrient stores and foragers with low nutrient stores is seen in numerous Hymenopteran species who evolved eusociality independently (Walton and Toth 2023). This convergent evolution has led to the hypothesis that nest worker-forager division of labor evolved through co-option of ancestral metabolic stress pathways (Walton and Toth 2023).

As central integrating signals in these metabolic stress pathways, neuropeptides are plausible targets of co-option, and recent studies implicate corazonin in this role: in *Harpegnathos saltator* (Jerdon, 1851) ants, corazonin promotes foraging behavior and inhibits the transition to the reproductive caste (Gospocic et al. 2017); in *Cataglyphis nodus* (Brulle, 1833) ants, the volume of corazonin-producing neurons was found to increase strongly as workers transition from in-nest tasks to foraging (Habenstein et al. 2021); while in honey bees (*Apis mellifera* Linnaeus, 1758), injection with synthetic corazonin led to increased foraging activity (Obernesser et al. 2025), although the downstream mechanism remains unclear. In *Harpegnathos* ants, corazonin appears to exert its effect at least partially by downregulating vitellogenin, the egg yolk protein found in nearly all oviparous animals. This control is of interest in honey bees (*Apis mellifera*), where vitellogenin and associated regulatory pathways were likely co-opted to influence the nest-worker to forager transition (Nelson et al. 2007). In addressing questions about whether and how metabolic stress pathways such as corazonin signaling were evolutionarily co-opted in eusocial insects, honey bees also provide a tractable system due to their experimental amenability, highly structured but plastic worker subcastes, and strong evidence for nutritional inequalities between nest workers and foragers.

This nutritional inequality is especially visible when comparing honey bee foragers with nurses, one of the main behavioral worker subcastes named for their primary role in feeding and caring for developing larval bees. Nurses have characteristically enlarged exocrine glands in the head (hypopharyngeal glands [HGs] and mandibular glands [MGs]) that they use to produce and secrete a nourishing, proteinaceous jelly that is fed to nestmates. To support this metabolically costly jelly production, nurses consume the most protein and lipid in the form of pollen. Compared to foragers, they have much higher fat body lipid stores (Toth and Robinson 2005), lipogenic capacity (Scofield and Amdam 2024), and protein synthesis, as well as higher vitellogenin synthesis and hemolymph titers (Piulachs et al. 2003; Nelson et al. 2007), resulting in higher starvation resistance (Fischer and Grozinger 2008). These differences are driven at least partially by differences in expression of numerous metabolic and nutritional signaling pathways including insulin/insulin-like growth factor signaling (IIS), ecdysteroid hormones, and neuropeptides (Ament et al. 2008; Ament et al. 2011b; Yamazaki et al. 2011; Han et al. 2015).

Preliminary evidence suggests that corazonin might have a role in regulating these differences: along with the pro-foraging behavioral role noted above, RNA-Seq data suggest differences in expression of *Crz* and *CrzR* between nurses and foragers (Gospocic et al. 2017; Harwood 2019). Given evidence for its role in eusocial and solitary insects, we hypothesized that corazonin is involved in regulating honey bee worker division of labor as a signal of metabolic stress. Specifically, we aimed to address three questions: 1) How do worker social role and acute metabolic challenge shape central and peripheral corazonin signaling? 2) Does corazonin regulate behavioral and physiological metrics associated with worker division of labor? 3) Is corazonin causally related to known molecular regulators of the nurse-forager transition, specifically vitellogenin, IIS, and ecdysteroids?

## Methods

### Collection of bees

Frames of sealed brood were collected from honey bee (*Apis mellifera ligustica*) colonies maintained at Arizona State University (Tempe, AZ, USA). Brood frames were left overnight in ventilated mesh boxes in a 34°C incubator humidified by placing an open container of water at the bottom of the incubator. Newly emerged workers were brushed from the three frames the next day and combined to form a mixed population to increase genetic diversity of the experimental groups. Bees were then used in subsequent experiments. All collections and handling procedures complied with institutional guidelines and relevant state and federal regulations governing honey bee research.

### Acute metabolic stress of nurses and foragers

To determine the effect of acute metabolic stress on corazonin signaling in honey bee workers, we followed our previous protocol for food restricting nurses and foragers of known age (Scofield and Amdam 2024). In brief, newly emerged bees were marked and released into host colonies and then collected on the same day: nurses were defined as 8-day old bees observed inserting their heads into larval cells for at least 3 seconds and foragers as 28-day old bees collected when returning to the hive entrance. Bees were placed into (5 x 12 x 16 cm) Plexiglass and mesh cages (30 bees/cage) and fed 30% sucrose solution in 12 mL syringes *ad libitum* for 3 hours. Cages were then either fed 30% sucrose solution for another 21 hours (feeding treatment) or instead fed only water for the same period (starvation), for a total of 24 hours of caging prior to sampling. Bees were collected from each of the four cages, anesthetized on ice, and their heads and fat bodies (complete abdominal cuticle with adhering fat body tissue minus the stinger, gut, crop, and ovaries) separately flash frozen in liquid nitrogen and maintained at -80°C until processed as described below. The food restriction experiment was carried out with bees sourced from separate colonies (two biological replicates for fat body qPCR, a single experimental cohort from one source colony for brain qPCR, and three biological replicate pools of ten brains per cage for peptide quantification). Sample sizes and technical details for each measurement are provided in the corresponding methodological sections below.

### RNA extraction and quantification of gene expression

To measure mRNA levels in the brain, flash frozen honey bee heads were dissected on dry ice to isolate the central complex of the brain, minus the optic lobes (referred to as “brains” throughout the text). Brains were then added to 350 µL of buffer RLT with a 0.5 mm steel bead, homogenized for 30s using a Retsch Mixer Mill MM 400, and then processed using a Qiagen RNeasy Mini Kit and Qiagen RNase-Free DNase according to manufacturer instructions. Fat body samples were processed similarly, except that they were homogenized in 1 mL of TRI reagent and RNA extracted using either a Qiagen RNeasy Mini Kit and Qiagen RNase-Free DNase or a Zymo Direct-zol Miniprep Kit and Zymo RNase-Free DNase I according to manufacturer instructions, including treatment with DNase I. RNA yield and quality were measured using a NanoDrop ND-1000 spectrophotometer, all samples from an experiment were diluted to the same concentration, and then stored at -80°C. Levels of mRNA in samples for genes of interest were quantified with real-time reverse transcription polymerase chain reaction (RT-qPCR) using QuantiTect SYBR Green RT-PCR kit (Qiagen) or ABScript II One Step SYBR Green RT-qPCR Kit (ABclonal) and ABI Prism 7500 (Applied Biosystems) as previously described (Amdam et al. 2004) using specific primers (Table S1). For the food restriction experiment, fat body *CrzR* expression was quantified across two biological replicates (*N* = 48 total, 6 bees per treatment group per colony), while brain *Crz* and *AKH* expression were quantified from a single experimental cohort from one source colony, with samples distributed across three technical qPCR plate runs that served as a blocking factor in statistical analyses ( *N* = 24 total). Gene expression was measured in comparison with *actin* by means of the delta-delta Ct method based on duplicate or triplicate readings.

### Liquid Chromatography–Tandem Mass Spectrometry (LC–MS/MS) analysis

To measure corazonin peptide levels, brains were dissected as described above, pooled in groups of 10 per cage, and homogenized in 250 µL of protein extraction buffer (8 M urea and 50 mM NH_4_HCO_3_) as in Gospocic et al. (2017) with 3 ng of recombinant corazonin (Vivitide) labeled with heavy phenylalanine (^13^C9, ^15^N, +10 Da) as an internal standard. Protein samples were prepared for LC-MS by subjecting them to solid phase extraction using Pierce C18 Spin Columns according to manufacturer instructions.

LC–MS/MS analyses were conducted on a Q Exactive Plus mass spectrometer (Thermo Fisher Scientific, San Jose, CA, USA) equipped with an EASY-Spray nano-electrospray ionization (nanoESI) source. Peptide samples containing both unlabeled corazonin peptide and isotopically labeled corazonin peptide incorporating heavy phenylalanine (13C_9, 15N) were first loaded onto an Acclaim PepMap 100 trap column (75 μm ID × 2 cm; Thermo Scientific) and subsequently separated on an Acclaim PepMap Neo UHPLC analytical column (75 μm ID × 50 cm; Thermo Scientific).

Chromatographic separation was achieved using a binary solvent system, where solvent A consisted of water with 0.1% formic acid and solvent B consisted of acetonitrile with 0.1% formic acid. Peptides were eluted using a gradient of 5–29% solvent B over 19 min, 29–99% over 5 min, followed by a column wash at 99% solvent B for 6 min. The flow rate was maintained at 300 nL/min using a Vanquish Neo UHPLC system (Thermo Fisher Scientific, San Jose, CA, USA).

Data-dependent acquisition (DDA) was performed using Xcalibur v4.7.69.37 software. Full MS survey scans were acquired at a resolution of 70,000 across an *m/z* range of 375–1600, with an automatic gain control (AGC) target of 1 × 10^6^ and a maximum injection time (IT) of 65 ms. The 14 most intense precursor ions were subjected to high-energy collisional dissociation (HCD) MS/MS with a normalized collision energy (NCE) of 27, a resolution of 17,500, an isolation window of 1.5 *m/z*, an AGC target of 5 × 10^4^, and a maximum IT of 65 ms. Ions with charge states of +1, 6, 7, 8, >8, unassigned, or isotopic peaks were excluded from fragmentation. A mass inclusion list containing precursor ions with charge states of +2, +3, +4, and +5 was employed to guide MS/MS acquisition.

Quantification of unlabeled and labeled corazonin peptides was based on the relative ratios of integrated peak areas corresponding to the extracted ion chromatograms at *m/z* 662.2917 and 667.3052, respectively.

Mass spectrometry was performed by the Office of Research, Innovation and Impact’s Analytical & Biological Mass Spectrometry Facility Core Facility at the University of Arizona (https://www.cores.arizona.edu/facilities/massspec).

### Synthetic corazonin and control injection

For injection, we used the only isoform of corazonin found in honey bees as treatment (Verleyen et al. 2006). A scrambled version of the peptide was used as the control (Table S2) following established approaches for neuropeptides (Choi et al. 2009). The control peptide sequence was selected from randomly generated sequences to have low sequence homology with corazonin so as not to induce a biological response. A previous study on corazonin in *Harpegnathos saltator* ants used injections of 100 ng of peptide dissolved in 0.5 µL of 1 × Hank’s balanced salt solution (Gospocic et al. 2017). Because the average wet weight of an *Apis mellifera* worker is approximately 110 mg, 3.44 times larger than the 32 mg average wet weight of an *H. saltator* worker, the amount of peptide injected was scaled up by that factor. Solutions for both corazonin and the scrambled control peptide were made up at concentrations of 350 ng/µL. In all experiments, bees were injected with 1 µL of the solution in the abdomen or thorax, with rationale for injection site described below for each experiment.

### Gustatory responsiveness and starvation resistance experiment

To test whether corazonin modulates short-term responses to metabolic stress, we standardized acute energetic state, upregulated corazonin via peptide injection, and measured changes in gustatory responsiveness and starvation resistance using standard protocols. We chose these metrics because they respond rapidly to changes in carbohydrate availability (Scheiner et al. 2001), differ across worker roles—with foragers showing higher and more labile gustatory responsiveness (Mayack et al. 2019)—and are sensitive to pheromonal, endocrine, and metabolic perturbation (Fischer and Grozinger 2008; Wang et al. 2012, 2016). Approximately 30 forager honey bees were collected from a colony, anesthetized on ice, and mounted into harnesses and secured with duct tape so that the proboscis could extend freely. All harnessed bees were fed to satiation with 30% sucrose solution to equalize their feeding status. Bees were then kept in a humidified box at room temperature for 18 hours, at which point they were fed 4 µL of 30% sucrose solution. Gustatory responsiveness was quantified using the proboscis extension response (PER) assay to produce a gustatory response score (GRS): each bee’s antennae were touched with ascending sucrose concentrations (0, 0.1, 0.3, 1, 3, 10, and 30%). To avoid sensitization, all bees were assayed in parallel with a 5-min interval between concentrations. GRS was calculated as the total number of proboscis extensions (Scheiner et al. 2001). Bees were then assigned to three groups: Crz and Scr groups that were injected with 350 ng of synthetic corazonin or scrambled corazonin, respectively, and noREF bees that were left untreated. Bees were injected into the thorax because access to the abdomen was obstructed by the harness. One hour after injection, GRS was measured again. After the GRS assay, the bees were placed back into the box and monitored for survival every 6 h for up to 72 h, following a previously published protocol with a longer sampling interval (Wang et al. 2012). All bees included in the starvation-resistance analysis died during the 72-h observation period; therefore, there were no right-censored observations. The entire experiment was replicated four times with bees from four different source colonies.

### Hemolymph sugar titer quantification

Hemolymph sugars were quantified in a separate experiment within the same acute energetic-state framework, focusing on glucose and trehalose as the primary circulating carbohydrates that reflect energetic state (Mayack et al. 2019). In this experiment, both nurses and foragers were collected, harnessed, fed, and injected as above to produce Crz or Scr groups. One hour after injection, 1 µL of hemolymph was collected from the median ocellus of each bee using a 1 µL glass microcapillary tube, added to a tube containing 299 µL of water, flash frozen in liquid nitrogen, and stored at -80°C until processing. This experiment comprised four independent experimental runs conducted with bees from three source colonies (replicates A and D originated from source colony A, while replicates B and C originated from colonies B and C, respectively). To quantify the glucose concentration, hemolymph samples were thawed, and 33 µL of sample and glucose standard curve added in triplicate to a 96 well plate along with 66 µL of Glucose (GO) Assay Kit Assay Reagent (Sigma Aldrich). Plates were sealed and incubated at 37°C for 30 min, 66 µL of sulfuric acid added, and then absorbance at 540 nm was quantified using a spectrophotometer. Glucose concentration in µg/µL was determined relative to the standard curve. Trehalose was measured similarly, except that the samples were first incubated at 37°C for 24 h with 1 U porcine trehalase. The amount of glucose produced from trehalose was calculated by subtracting the first reading from the second reading. The amount of trehalose was estimated using the equation: trehalose (μg)=glucose (µg)×342.3/(180.2×2) (Hartfelder et al. 2013; Wang et al. 2013).

### Corazonin injection and gene expression experiments

To test whether corazonin affects fat body gene signaling, nurse honey bees were collected from colonies, injected as above to produce Crz and Scr groups, and then maintained in cup cages in an incubator at 31°C. Twenty-four hours after injection, fat bodies were sampled from bees as described above. The experiment was replicated twice using two different source hives. In a separate experiment, 7-day old bees were treated as Crz, Scr, or noREF groups above, and returned to a host hive. Twenty-four hours later, bees were recovered, anesthetized on ice, and 1 µL of hemolymph was collected using a glass capillary tube after puncturing the abdomen between the second and third tergites with a sterile BD PrecisionGlide G30 x 1/2 needle. Care was taken to avoid contaminating the samples with tissue fragments and foregut content. Hemolymph (1 µL) was dissolved in 10 µL phosphate-buffered saline with a cOmplete protease inhibitor (Sigma-Aldrich) and stored at -20°C until analysis. The fat body of each worker was then dissected, homogenized in 500 µL of TRIzol (Invitrogen, Carlsbad) and stored at -80°C until RNA could be extracted.

### Preparation of dsRNA

A custom computational script was used to select a 500bp region of the honey bee *Corazonin receptor* (*CrzR,* NM_001143921.1) and vitellogenin gene (*Vg*, NM_001011578.1) with no 20-bp and minimal 19-bp homologies to the rest of the honey bee transcriptome (plasmid sequences in supplemental). TRIzol extraction and reverse transcription were used to produce cDNA from honey bee fat body tissue according to standard procedures. Phusion High-Fidelity DNA polymerase (New England BioLabs) was used with the specific primers (Table S3) to produce the selected sequence in two different orientations with a 16-bp 5’ overhang needed for ligation using the In-Fusion HD Cloning system (Takara Bio): PCR products were purified using the NucleoSpin PCR purification kit, before being directionally inserted into the MCS of the pBluescript SK (+) vector downstream of the T7 promoter. Bacterial transformation was accomplished according to standard protocols, before using a QIAGEN Plasmid Midi Kit to isolate the plasmids from clonally selected cells. The sequence of the plasmids was verified by Sanger sequencing and aligned against the honey bee genome. Linearized plasmid was used as template for transcription reactions using the Promega RiboMax T7 system to produce single-stranded RNA for sense and antisense strands of the *CrzR* and *Vg* clones, which were then combined at 70°C and allowed to cool to room temperature. The resulting dsRNA was purified using TRIzol reagent before renaturation and suspension in nuclease-free water at a concentration of 10 µg/µL.

### Double knockdown experiment

As an additional approach to test for regulatory interactions between corazonin and vitellogenin, we used RNA interference targeting *Vg* and *CrzR*. Frames containing large amounts of capped brood were retrieved from three different *Apis mellifera* source colonies maintained at the ASU Honey Bee Research Lab and placed in an incubator overnight at 34°C. Newly-emerged workers were randomly assigned to one of four treatments: (1) noREF, a non-injected reference group; (2) injC, a control group injected with vehicle (nuclease-free H_2_O); (3) *CrzR-*RNAi, a dsRNA-injected *CrzR* knockdown group; and (4) *Vg-*RNAi, a dsRNA-injected *Vg*-knockdown group. Workers from all treatment groups were anesthetized on ice, immobilized on a clean wax disk with two pins, and marked with paint to indicate treatment identity. Injections were performed between the fifth and sixth tergites using Hamilton syringes with G30 disposable needles. Injection volume was 1 µL. Bees showing signs of hemolymph leakage after withdrawal of the needle were discarded. After treatment, bees were placed in a host colony. Seven days later, workers were recovered, anesthetized on ice, and 1 µL of hemolymph was collected using a glass capillary tube after puncturing the abdomen between the second and third tergites with a sterile BD PrecisionGlide G30 x 1/2 needle. Care was taken to avoid contaminating the samples with tissue fragments and foregut content. Hemolymph (1 µL) was dissolved in 10 µL phosphate-buffered saline with a cOmplete protease inhibitor (Roche, Basel, Switzerland) and stored at-20°C until analysis. The fat body of each worker was then dissected, homogenized in 500 µL of TRIzol (Invitrogen, Carlsbad, CA, USA) and stored at -80°C until RNA could be extracted. To avoid potential false-positive amplification of residual injected dsRNA during RT-qPCR knockdown verification, qRT-PCR measurement primers for *CrzR* (CrzR-7; Table S1) were specifically designed across exon junction 2/3 outside the 500-bp dsRNA target region, ensuring that only endogenous mRNA was amplified.

### Hemolymph vitellogenin titer quantification

Individual hemolymph samples were separated by one-dimensional SDS electrophoresis using 7.5% polyacrylamide gels. Each gel included two samples from each treatment group, one known protein marker (Bio-Rad), and three standard samples (1.25, 2.5, and 3.75 µg) of *Escherichia coli* β-galactosidase (Sigma), randomly distributed among lanes. Sample aliquots containing 0.2 µL of hemolymph were mixed with loading buffer (Laemmli, 2×, Sigma) and heated at 95°C for ten minutes. Electrophoresis was carried out according to published protocols (Lin et al. 1999; Amdam et al. 2004). The analysis was performed using ImageJ after staining the gels with Coomassie Brilliant Blue (Sigma-Aldrich). The vitellogenin band (180–190 kDa) was quantified relative to the β-galactosidase protein standard.

### Statistical analysis

Statistical analyses were performed in R using the model or test specified for each experiment. Brain *Crz* and *AKH* expression (*N* = 24) were analyzed using Type II ANOVA with worker class, feeding treatment, and brain qPCR plate as predictors. Type II ANOVA was also used for the peptide, gustatory responsiveness, glucose, and trehalose models, with the corresponding experimental factors and blocking variables. For hemolymph glucose and trehalose titers, linear models evaluated caste, injection treatment, and their interaction, with experimental run (replicate) included as a blocking factor (response ∼ caste * treatment + replicate). Because runs were nested within source colonies (colonies B and C each contributed a single run, and colony A contributed two), source-colony effects were confounded with run and could not be independently estimated. Ordinary ANOVA models were used for the starvation-resistance analysis, the Fig. 4 gene-expression analyses, and the one-way Fig. 5 pilot analyses. The Fig. 5C comparisons were followed by Tukey’s HSD.

For fat body *CrzR* expression in the starvation experiment, a Kruskal-Wallis test across the four treatment groups was followed by Dunn’s test with a Bonferroni correction for the family-wise error rate (FWER). The factorial interaction between worker class and feeding treatment was tested using an aligned rank transform (ART) model implemented with the art function in the R package ARTool (Kay et al. 2025), with worker class and feeding treatment as fixed factors and experimental replicate as a random blocking factor (CrzR ∼ class * treatment + (1|replicate)). For the RNAi experiments, log10-transformed responses were analyzed with factorial ANOVA: Fig. 6A used treatment × replicate × plate, Fig. 6B used group × replicate followed by pooled treatment-only Tukey’s HSD, and Fig. 6C used treatment × plate.

To evaluate the relationship between relative fat body *Vg* mRNA expression and circulating hemolymph vitellogenin protein titer in the peptide injection pilot trial (*N* = 60), an ordinary least-squares linear regression model was fit using the base R lm function, and the coefficient of determination (*r*^2^), *F*-statistic, and *P*-value were calculated (Fig. S1).

## Results

### Corazonin signaling differs between nurses and foragers

To look for evidence that corazonin signaling is involved in division of labor and caste-specific metabolic regulation in worker honey bees, we used a two-factor experiment where 8-day-old nurses and 28-day-old foragers were either fed 30% sucrose solution *ad libitum* or starved (given only water) for 21 hours in cages. Brain levels of *Crz* mRNA were significantly higher in foragers than in nurses (Fig. 1A; *F*_1,18_ = 28.14, *P* < 0.001) and not affected by starvation (*F*_1,18_ = 0.47, *P* = 0.50). There was no significant effect of brain qPCR plate (*F*_2,18_ = 1.07, *P* = 0.36) or of the interaction between worker caste and starvation (*F*_1,18_ = 0.33, *P* = 0.57). Because AKH is a closely related metabolic peptide in the same neuroendocrine family, we also measured AKH transcript levels to determine whether any changes in corazonin signaling reflected a broader shift in neuropeptide expression. Brain levels of *AKH* mRNA were not affected by worker caste (Fig. 1B; *F*_1,18_ = 0.57, *P* = 0.46), starvation (*F*_1,18_ = 0.42, *P* = 0.53), brain qPCR plate (*F*_2,18_ = 2.06, *P* = 0.16), or the interaction between worker caste and starvation (*F*_1,18_ = 0.13, *P* = 0.73). The quantity of corazonin peptide in forager brains was significantly higher than in nurses (Fig. 1C; *F*_1,8_ = 68.97, *P* < 0.001) and not affected by starvation (*F*_1,8_ = 0.42, *P* = 0.53) or caste × starvation (*F*_1,8_ = 0.00, *P* = 0.99). Fat body levels of *CrzR* mRNA differed significantly across the four treatment groups (Fig. 1D; Kruskal-Wallis, *χ*^2^(3) = 12.35, *P* < 0.01), and an aligned rank transform analysis showed a significant interaction between worker caste and starvation (ART, *F*_1,43_ = 6.31, *P* = 0.016). A post hoc Dunn’s test with a Bonferroni correction for FWER showed that starved nurses had significantly higher *CrzR* expression relative to fed nurses (*P* < 0.01); there were no other significant differences between groups. Taken together, these results suggest that central corazonin signaling is shaped primarily by worker caste rather than acute metabolic challenge, whereas the significant worker caste × starvation interaction in fat body *CrzR*, driven by starvation-associated upregulation in nurses but not foragers, supports a caste-specific response to starvation.

**Fig. 1.**
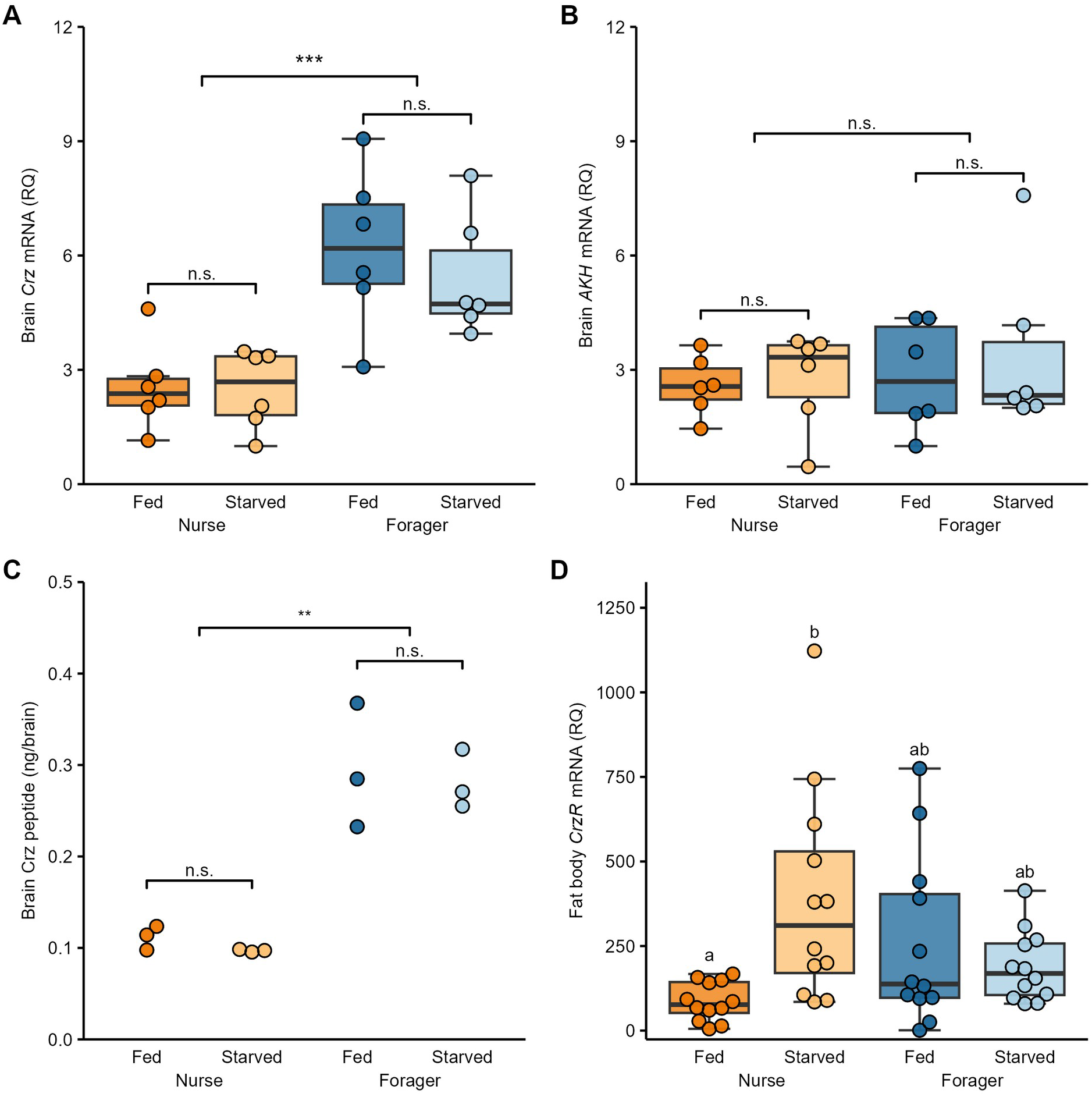
Crz signaling is higher in foragers and in starved nurses. (A) Foragers have significantly higher brain *Crz* gene expression than nurses while starvation has no effect. (B) Brain *AKH* expression is not affected by worker caste or starvation. (C) Forager brains have significantly higher quantities of corazonin peptide than those of nurses while starvation has no effect. (D) Starvation significantly increases the fat body expression of *CrzR* in nurses but not in foragers. Points show individuals (A, B, D) or pooled samples of 10 brains (C); boxplots show IQR, median, and range.

### Corazonin injection had no detected effect on gustatory responsiveness, starvation resistance, or hemolymph sugar

To further assess whether corazonin signaling mediates the acute response to metabolic stress, we measured gustatory responsiveness (GRS), starvation resistance, and hemolymph glucose and trehalose titers, which are standard metrics of energetic state. If corazonin modulates carbohydrate metabolism in honey bees as it does in *Drosophila* (Kubrak et al. 2016), we reasoned that upregulation of corazonin might alter these metrics. We therefore injected forager honey bees with 350 ng of corazonin or a scrambled control peptide (Scr). An additional group of bees was left untreated (noREF). Corazonin treatment had no significant effect on the difference between GRS measured before and 1 hour after injection (Fig. 2A; *F*_2,134_ = 0.45, *P* = 0.64) or on starvation resistance (Fig. 2B; *F*_2,79_ = 0.75, *P* = 0.48). In a second injection experiment, corazonin injection did not affect hemolymph glucose (Fig. 3A; *F*_1,50_ = 1.12, *P* = 0.29) or trehalose titers (Fig. 3B; *F*_1,50_ = 0.0007, *P* = 0.98) 1 hour after injection in either nurse or forager honey bees. Foragers had significantly higher glucose (*F*_1,50_ = 5.58, *P* = 0.02) and trehalose titers (*F*_1,50_ = 27.21, *P* < 0.001) than nurses; there was a significant replicate effect for glucose (*F*_3,50_ = 7.77, *P* < 0.001) but not for trehalose titers (*F*_3,50_ = 2.03, *P* = 0.12). There was no significant caste × treatment interaction for either glucose (*F*_1,50_ = 0.75, *P* = 0.39) or trehalose titers (*F*_1,50_ = 0.20, *P* = 0.66). We thus detected no effect of corazonin on these carbohydrate-related metrics under the tested dose and timeframe.

**Fig. 2.**
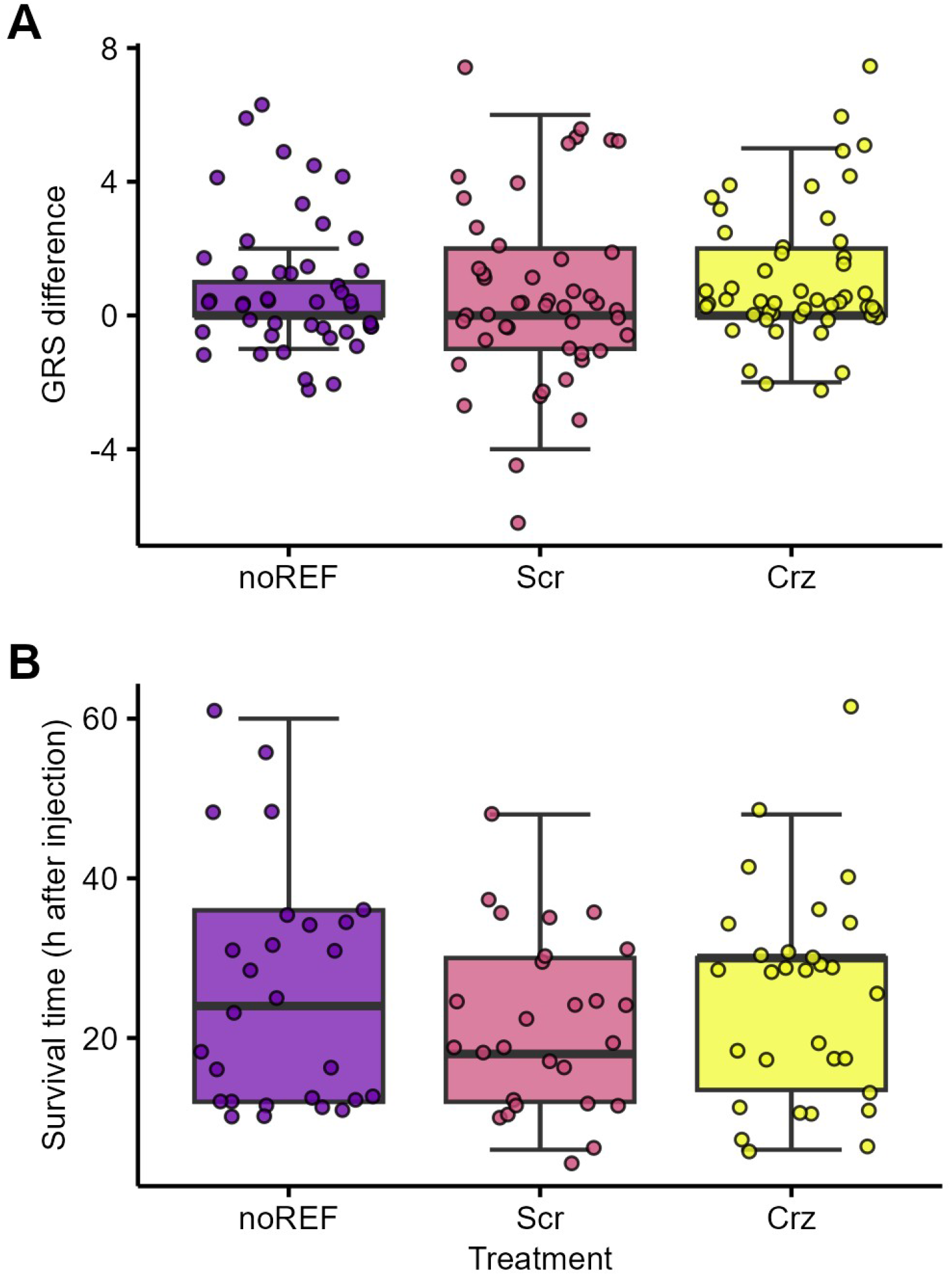
Crz injection does not affect gustatory responsiveness or starvation resistance. (A) GRS is not affected by injection. (B) Survival time under starvation conditions is not affected by the injection treatments.

**Fig. 3.**
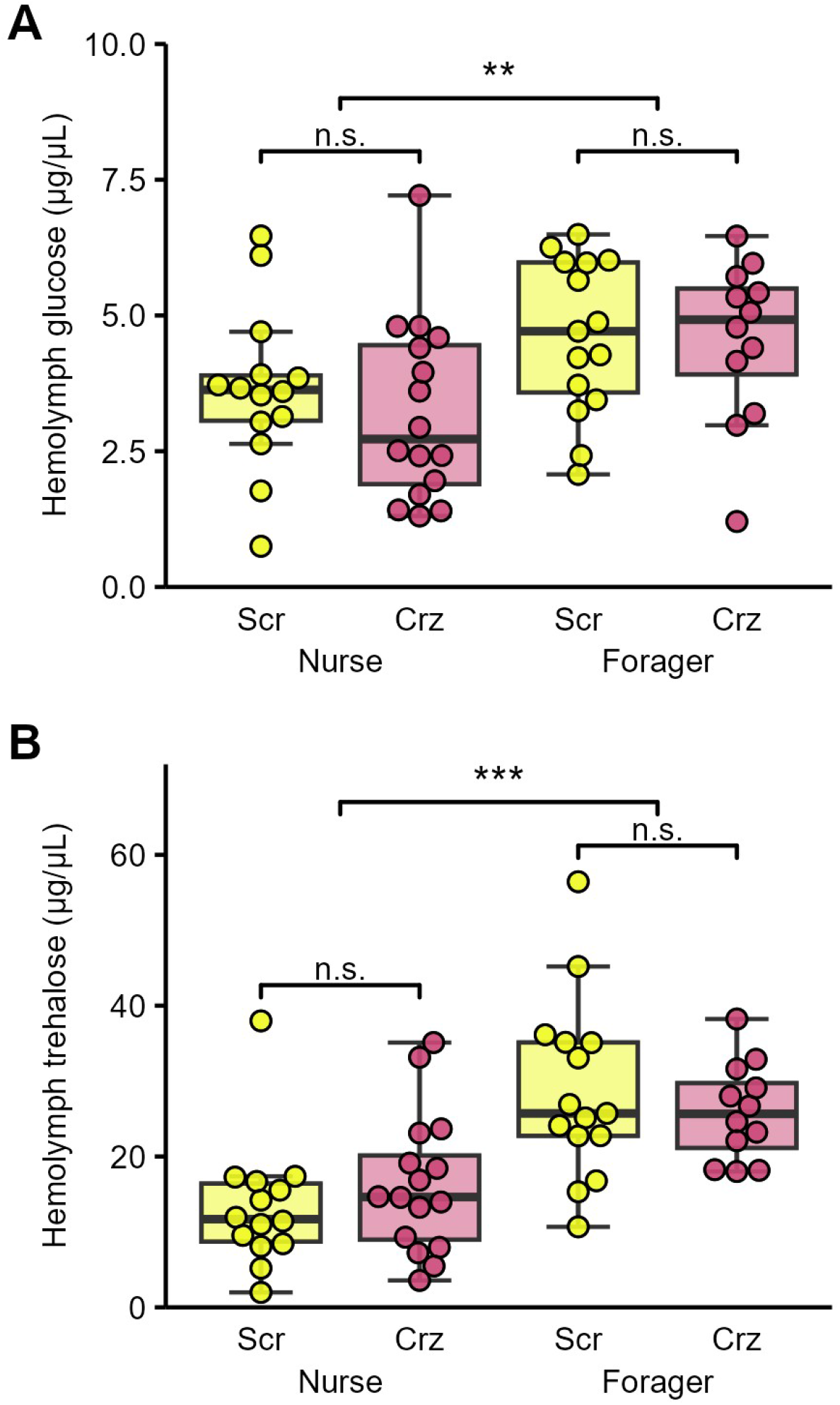
Crz injection does not affect hemolymph sugar (A) Hemolymph glucose titers are not affected by injection in either nurses or foragers, but foragers have significantly higher glucose titers than nurses. (B) Similarly, hemolymph trehalose titers are not affected by injection, but foragers have significantly higher trehalose titers than nurses.

### Corazonin injection downregulates IRS transcript levels in worker fat body

To look for evidence whether corazonin signaling influences other endocrine signaling pathways, we measured transcripts of *IRS*, *EcRB*, *UpdlA*, and *Cyp314a1* following peptide injection. These genes represent distinct components of IIS, ecdysteroid, and adipokine signaling linked to corazonin in other insects, providing a broad survey of pathways potentially responsive to corazonin. Injection of nurse honey bees with 350 ng of synthetic corazonin significantly lowered fat body *IRS* transcript levels relative to injection with a scrambled control peptide (Fig. 4A; *F*_1,12_ = 5.41, *P* = 0.038). Replicate (*F*_1,12_ = 0.29, *P* = 0.60) and the interaction between replicate and injection (*F*_1,12_ = 0.01, *P* = 0.91) were not significant. Injection did not affect transcript levels of *EcRB* (Fig. 4B; *F*_1,12_ = 0.16, *P* = 0.70), *UpdlA* (Fig. 4C; *F*_1,12_ = 0.06, *P* = 0.81), or *Cyp314a1* (Fig. 4D; *F*_1,12_ = 1.79, *P* = 0.21). There were no significant replicate effects for *EcRB* (*F*_1,12_ = 3.33, *P* = 0.09), *UpdlA* (*F*_1,12_ = 0.70, *P* = 0.42), or *Cyp314a1* (*F*_1,12_ = 1.77, *P* = 0.21). Likewise, there were no significant interactions between replicate and injection for *EcRB* (*F*_1,12_ = 1.09, *P* = 0.32), *UpdlA* (*F*_1,12_ = 0.66, *P* = 0.43), or *Cyp314a1* (*F*_1,12_ = 0.01, *P* = 0.92). The specific downregulation of *IRS* transcript levels without affecting other pathway gene transcripts suggests an interaction between corazonin and IIS.

**Fig. 4.**
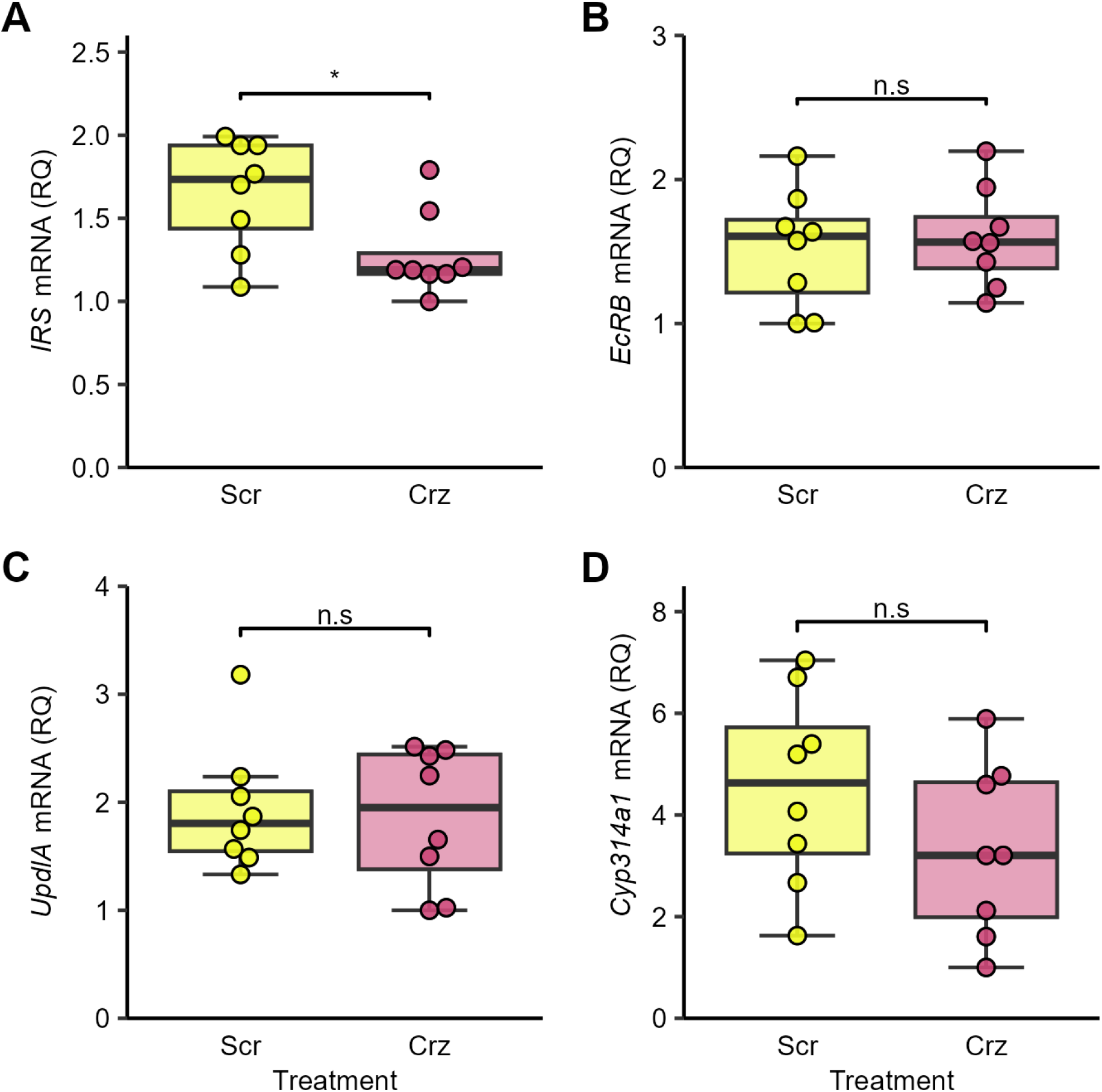
Crz injection downregulates fat body insulin receptor substrate (*IRS*) expression. (A) Nurse worker bees injected with 350 ng of synthetic corazonin (Crz) have significantly lower abdominal insulin receptor substrate (IRS) expression 24 hours after injection than those injected with a scrambled control (Scr). (B) *Ecdysone Receptor B* (*EcRB*), (C) *Unpaired-like A* (*UpdlA*), and (D) *Cyp314a1* are not affected.

### No evidence for a corazonin-vitellogenin regulatory axis

In a separate set of experiments, we explored a previously hypothesized corazonin-vitellogenin regulatory axis (Gospocic et al. 2017). Given the centrality of vitellogenin in honey bee worker behavior, we explored both directions of causation, first upregulating corazonin signaling by injection, and second downregulating *Vg* expression by RNAi. We also attempted to downregulate corazonin signaling by RNAi targeting *CrzR* but this had low efficacy and thus functioned as an RNAi control. Injection of 7-day-old bees with 350 ng of synthetic corazonin had no significant effect on *Vg* fat body gene expression (Fig. 5A; *F*_2,57_ = 0.73, *P* = 0.49) or hemolymph vitellogenin titer (Fig. 5B; *F*_2,57_ = 0.54, *P* = 0.59) 24 hours after injection. There was a significant relationship between *Vg* expression and vitellogenin titer (Fig. S1; *r*^2^ = 0.2998, *F*_1,58_ = 24.83, *P* < 0.001). Fat body expression of *CrzR* was significantly affected by treatment (Fig. 5C; *F*_2,57_ = 3.84, *P* = 0.03), with post hoc testing showing that the corazonin-injected bees had significantly higher *CrzR* expression than the noREF bees (*P* = 0.04). In the RNAi experiment targeting *Vg* and *CrzR*, fat body gene expression of *Vg* was significantly affected by treatment (Fig. 6A; *F*_3,32_ = 15.08, *P* < 0.001), replicate (*F*_1,32_ = 27.54, *P* < 0.001), and qPCR plate (*F*_1,32_ = 0.02, *P* = 0.90). Post hoc testing showed that the Vg-RNAi group had significantly lower *Vg* expression than the noREF (*P* < 0.001), injC (*P* < 0.01), and CrzR-RNAi groups (*P* < 0.01). Hemolymph vitellogenin titer was significantly affected by treatment (Fig. 6B; *F*_3,40_ = 4.52, *P* = 0.008) and replicate (*F*_1,40_ = 12.71, *P* = 0.001). Post hoc testing of the pooled data showed that the Vg-RNAi group had significantly lower vitellogenin titer than the noREF group (*P* = 0.014), but not the injC group (*P* = 0.079). *CrzR* gene expression was not affected by treatment (Fig. 6C; *F*_3,16_ = 0.41, *P* = 0.75) or qPCR plate (*F*_1,16_ = 0.17, *P* = 0.69). We thus found no effect of corazonin on vitellogenin under the tested dose and timeframe.

**Fig. 5.**
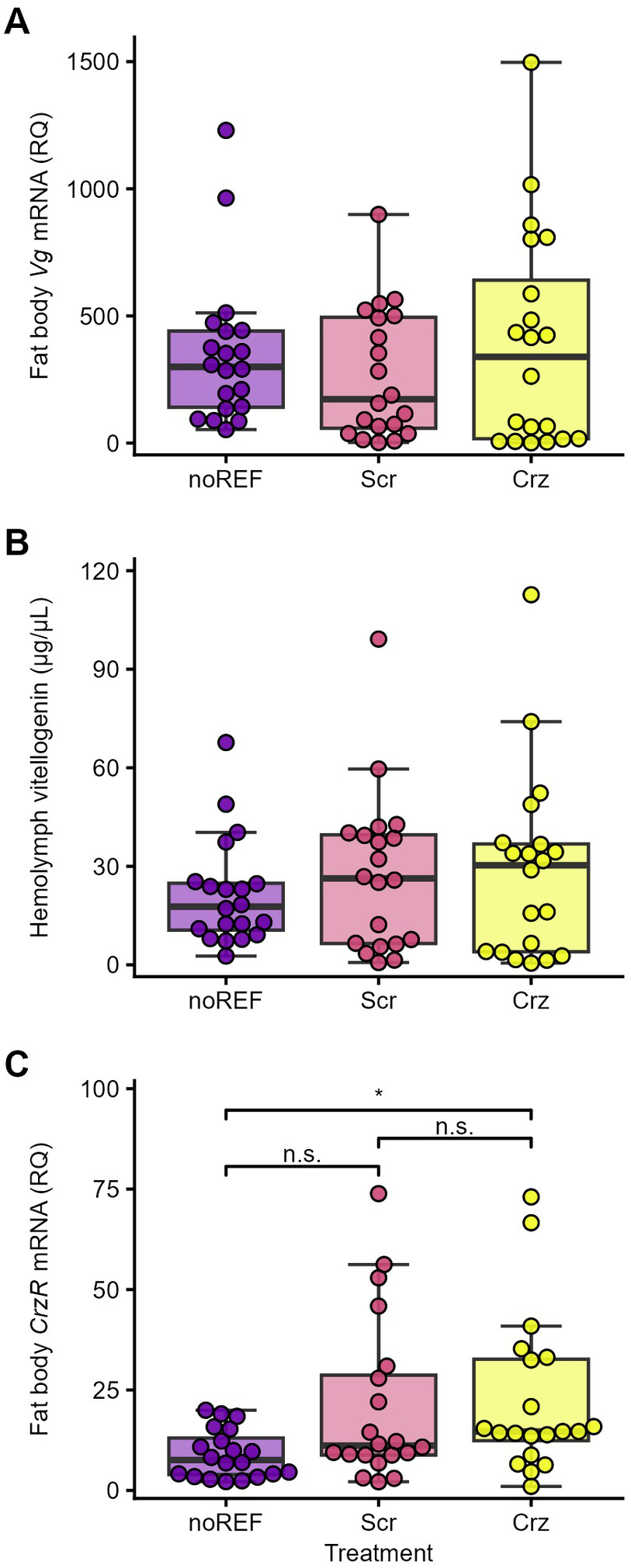
Crz injection does not suppress vitellogenin synthesis. Seven-day old worker bees injected with 350 ng of synthetic corazonin (Crz) do not have significantly different (A) fat body *Vg* expression or (B) hemolymph vitellogenin titers 24 hours after injection than those injected with a scrambled control (Scr) or not injected (noREF). (C) Fat body *CrzR* expression is significantly higher in Crz-injected bees than in controls.

**Fig. 6.**
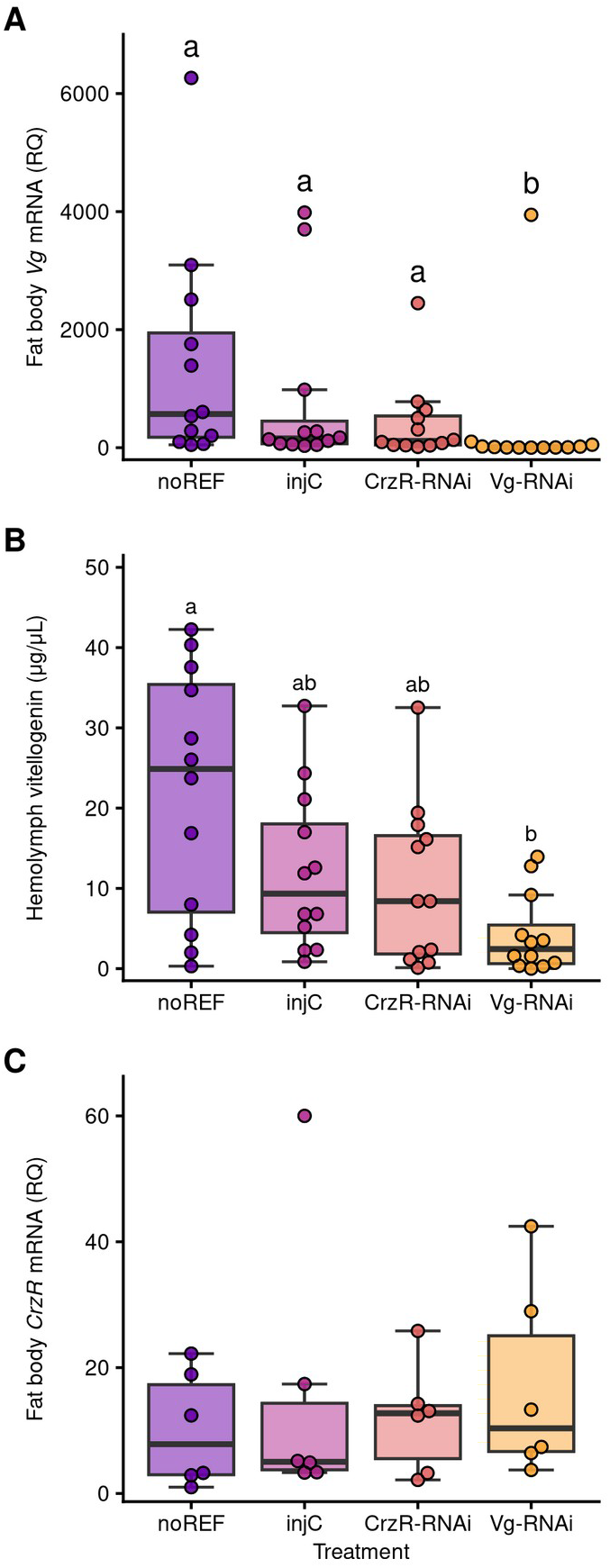
Knockdown of Vg does not affect CrzR expression. (A) Injection of 7-day old bees with dsRNA against *Vg* significantly lowers fat body expression relative to bees injected with vehicle (injC) or dsRNA against *CrzR* or bees not injected (noREF). (B) *Vg* RNAi causes significantly lower hemolymph vitellogenin titer compared with noREF bees; the comparison with injC bees is not significant. (C) Fat body *CrzR* expression is unaffected by treatment.

## Discussion

In this study, we found that corazonin signaling differs between worker castes, with central corazonin upregulation in foragers and starvation-induced corazonin responsiveness in the fat body of nurses. Complementary molecular experiments further revealed a targeted effect of corazonin on insulin receptor substrate (*IRS*) expression and indicated specificity relative to other major nutritional and endocrine pathways.

By measuring both gene expression and peptide levels in the brains of fed and starved nurses and foragers, we showed that foragers have approximately three-fold higher brain corazonin levels than nurses (Fig. 1A, C). This pattern aligns with past work linking corazonin brain levels to foraging activity in ants and supports the idea that corazonin is a conserved marker of foraging behavior. For example, in *Cataglyphis* ants, the volume of corazonin-producing neurons in the brains of workers increases along with *Crz* gene expression as workers transition from inside-nest tasks to foraging, although corazonin peptide levels were not measured directly (Habenstein et al. 2021). In *Harpegnathos* ants, corazonin peptide levels are higher in the brains of foragers than in egg-laying gamergates, but the study did not compare levels among different worker subcastes (Gospocic et al. 2017). To our knowledge, our study is the first to use mass spectrometry to quantify corazonin peptide levels directly in workers with different social roles. Interpreting what these levels mean requires considering the balance among production, storage, release, and degradation. Although peptide levels themselves cannot specify which process has changed, the matched three-fold increase in corazonin peptide (Fig. 1C) and *Crz* transcript levels in foragers (Fig. 1A), together with the absence of a starvation effect on either measure, are most consistent with a forager-linked increase in synthesis and maintenance of a larger releasable pool rather than an acute, starvation-driven change in release. Moreover, despite multi-year attempts using a sensitive LC-MS workflow, we were unable to detect corazonin in worker hemolymph across diverse contexts, whereas brain peptide was readily quantifiable. This could be because the corazonin signaling is primarily central rather than peripheral (though the strong *CrzR* upregulation we measured in starved nurses argues in favor of at least some peripheral signaling), or because the amount of peptide released is relatively small. Our approach is also unable to tell us anything about release dynamics, such as the pulse frequency dependent effects seen in the homologous peptide hormone GnRH, where a change in frequency of release causes different physiological outcomes (Burger et al. 2002; Thompson and Kaiser 2015). Moderate differences in release thus remain possible and will require direct assays to resolve (e.g., imaging of corazonin-producing neurons).

We explored one possible function of corazonin as a glucagon-like signal that could be released in response to low hemolymph sugar to mobilize stored nutrients. This function is supported in *Drosophila* male flies, where knockdown of *CrzR* in the fat body increases the flies’ resistance to starvation stress and elevates circulating and stored carbohydrates (Kubrak et al. 2016). Corazonin signaling has a diversity of functions across insects, and here, we did not detect such an effect under the tested dose and timeframe. Starvation did not significantly affect brain *Crz* gene expression (Fig. 1A) or corazonin peptide levels (Fig. 1C) in nurses or foragers, suggesting that acute starvation stress does not cause an increase in central-to-peripheral corazonin signaling. Consistent with this lack of response, we detected no effect of synthetic corazonin injection on GRS (Fig. 2A), starvation resistance (Fig. 2B), or hemolymph glucose (Fig. 3A) or trehalose (Fig. 3B) under the tested dose and timeframe. These factors are known to be co-regulated: for example, joint knockdown of *Vg* and the juvenile-hormone-response gene *ultraspiracle* increases GRS, lowers starvation resistance, and increases hemolymph glucose and trehalose but does not affect lipid metabolism (Wang et al. 2012). Together with the absence of evidence for increased corazonin release during starvation, we detected no effect of corazonin on the carbohydrate metabolism regulatory module under the tested dose and timeframe.

Intriguingly, however, fat body *CrzR* expression was significantly increased by starvation in nurses but not in foragers, with starved nurses having an approximately 4.5-fold increase in *CrzR* expression relative to fed nurses (Fig. 1D). A possible explanation for this result is that fat body sensitivity to starvation-induced corazonin signaling is state-dependent: during starvation, nurse fat bodies become more sensitive to corazonin signaling through upregulating CrzR, while forager fat body corazonin sensitivity appears to remain static, or at least lack a transcriptionally regulated increase. The lack of starvation-induced *CrzR* expression in foragers could mean that foragers have a reduction in peripheral corazonin sensitivity; alternatively, forager corazonin signal transduction could already be at a ceiling level, meaning no increase in *CrzR* expression is needed. Since we did not directly measure CrzR levels and correlation between transcript and protein levels is often poor (Wegler et al. 2020), our data cannot distinguish between these possibilities. This is especially relevant for GPCRs like CrzR, where signal transduction is regulated through mechanisms independent of mRNA levels, such as phosphorylation, spatial organization, and endocytosis (Weinberg and Puthenveedu 2019). Regardless, whether forager corazonin signaling is at a floor or ceiling, the state-dependent sensitivity in transcriptional upregulation is consistent with other developmental checkpoints: an example is found in ovarian development in *Drosophila*, where ovariole number is much more sensitive to poor nutrition before reaching a “critical weight” developmental checkpoint than after (Mendes and Mirth 2016). The switch in sensitivity results from a reprogramming of tissue-level sensitivity to hormonal signals, resulting in developmental stages with different responses to nutritional state.

Similarly, honey bee nurses and foragers can be viewed as distinct developmental stages, separated by a highly plastic developmental checkpoint that responds to many nutritional and non-nutritional signals. In this framework, nurses are the high-nutrition developmental stage, which is reflected in their much larger and more metabolically active fat body compared to foragers (Haydak 1957; Toth and Robinson 2005; Ament et al. 2011a; Chan et al. 2011; Scofield and Amdam 2024). The lower-nutrition forager stage is not a general starvation phenotype, but rather reflects differential allocation of specific nutritional resources within the colony. Maintenance of the nurse metabolic state depends on pollen-derived protein and lipid to support brood rearing, making pollen-derived nutrition a limiting factor for colony growth (Arien et al. 2020; Stabler et al. 2021). Recent evidence indicates that social displacement of aging workers from the pollen-rich brood area reduces their access to pollen, and that this nutritional shift drives the transition from nursing to foraging (Zanni et al. 2026). Consistent with this nutritional regulation of the transition, nurses retain developmental plasticity, remaining competent to produce an appropriate developmental state in response to nutritional status. Foragers have lost this plasticity: while under certain conditions, such as colony-level need for nurses, foragers can revert to nursing, this reversion cannot be induced by nutritional cues; i.e., while low nutrition can induce the nurse-to-forager transition, high nutrition cannot induce the forager-to-nurse transition (Schulz et al. 1998; Toth et al. 2005). Starvation-induced fat body *CrzR* expression in nurses but not in foragers is consistent with this canalization of foragers into a permanent low-nutrition state with a corresponding reduction in responsiveness to metabolic stress, and this pattern suggests that corazonin signaling is involved in regulating metabolic plasticity in workers.

We found a possible downstream target of this corazonin signal in our injection experiment, where injecting nurses with corazonin caused downregulation of the IIS pathway gene *IRS* (Fig. 4A) but not *UpdlA* or the ecdysteroid genes *EcRB* or *Cyp314a1* (Fig. 4B–D). The IIS pathway is known to play a central role in division of labor in honey bees: during larval development, several IIS pathway genes are expressed at much higher levels in queen larvae than in worker larvae (Wheeler et al. 2006; Azevedo and Hartfelder 2008). Since the primary trigger determining whether a larva becomes a queen or a worker is quantity and quality of the jelly it is fed (Patel et al. 2007), these differences in IIS are likely diet-induced, with high nutrition producing the queen phenotype through a regulatory network that involves IIS pathway genes.

IIS also contributes to adult worker division of labor in a manner consistent with nutrition-mediated phenotypic plasticity. Foragers show higher brain *ILP1* and abdomen *InR1* and *InR2* than nurses, and pollen deprivation induces a forager-like expression pattern (Ament et al. 2008). In diet experiments, expression of *ILP1* was affected by diet composition and was highest on a diet high in protein, while expression of *ILP2* and *IRS* was not affected by diet, supporting a role for the IIS pathway in integrating nutritional inputs (Ihle et al. 2014). Concurrently, injection of bees with amino acids increased *ILP1* expression (Nilsen et al. 2011). In behavioral experiments, knockdown of *IRS* (likely primarily in the fat body) affected foraging behavior, with *IRS* knockdowns collecting less nectar and more pollen but having similar GRS compared with controls (Wang et al. 2010). *IRS* knockdowns also lived shorter lives and foraged earlier (Ihle et al. 2019). Treatment of summer nurse bees with bovine insulin significantly decreased the age at which foraging was initiated (Mott and Breed 2012). These results support the hypothesis that IIS is involved in mediating food-seeking behavior in response to nutritional stress.

Corazonin has similarly already been linked to food-seeking behavior in both social and non-social insects: in *Drosophila*, reduction of corazonin signaling by knockdown of peripheral (fat body and salivary gland) *CrzR* caused a reduction in feeding behavior (Kubrak et al. 2016), while an increase in corazonin signaling by injection with corazonin promoted foraging behavior in *Harpegnathos* ants (Gospocic et al. 2017) and honey bee workers (Obernesser et al. 2025). The mechanism by which corazonin modulates behavior (especially via peripheral signaling) is not fully understood, but there is support for IIS as a downstream target: along with our finding in this study that corazonin injection downregulates *IRS* in honey bee fat body, work in *Drosophila* shows that peripheral *CrzR* knockdown affects expression of insulin-like peptides in the brain and insulin-regulated genes in the fat body (Kubrak et al. 2016). Additionally, we recently found a very strong (*r* = 0.86) correlation between fat body *CrzR* and *IRS* gene expression in 8-day-old workers in a cage experiment along with both genes being upregulated by protein deprivation, further linking corazonin and IIS as fat body regulatory elements responsive to the same nutritional cues (Scofield and Amdam 2026). Corazonin functioning to modulate IIS would explain the lack of acute response to corazonin injection found in this and previous studies and give a mechanism by which chronic behavioral and developmental changes are induced, such as the increase in foraging activity found in corazonin-injected bees from 3 to 6 days post-treatment (Obernesser et al. 2025). However, while this is plausible, it is important to note that injection studies like these that globally upregulate corazonin cannot determine the site at which the effect is produced. Given the expression of *CrzR* in brain and fat body, it is unclear whether the reduction in fat body *IRS* levels we found in this study is produced by effects on brain ILP synthesis or release, by direct action on the fat body, or by some other unknown mechanism. The inefficacy of fat body *CrzR* knockdown in this study makes RNAi a less promising approach. Further work is needed to map the tissue-specific role of corazonin signaling in honey bees.

Besides IIS, a second proposed downstream target of corazonin signaling is vitellogenin: in *Harpegnathos* ants, *Crz* expression is high in foragers and low in reproductive, nest-confined gamergates, and injection of synthetic corazonin peptide causes downregulation of *Vg* expression and increased foraging behavior (Gospocic et al. 2017). We found no evidence of a corazonin-vitellogenin relationship in honey bee workers: injection with synthetic corazonin had no effect on fat body *Vg* expression (Fig. 5A) or hemolymph vitellogenin titers (Fig. 5B), while knockdown of *Vg* did not affect fat body *CrzR* expression (Fig. 6C). While the negative result should be interpreted cautiously, it suggests that the conserved function of corazonin as a pro-foraging signal is not dependent on the corazonin-vitellogenin relationship found in *Harpegnathos*, although further comparative and experimental work is needed to understand the dynamics of the corazonin-IIS link found in our study and to determine if it is present across the Hymenoptera.

In conclusion, our findings clarify the role of corazonin in honey bee worker biology. At the level of direct experimental evidence, we demonstrate that age-typical foragers have more brain *Crz* transcript and peptide, that starvation increases fat-body *CrzR* transcript in nurses, and that injected corazonin reduces *IRS* transcript under the tested conditions. Together with previous work, these findings connect corazonin signaling with IIS and the nutritional regulation of worker state, though the regulatory architecture, tissue-specific mechanisms, and physiological consequences of this relationship remain unresolved. At the broadest evolutionary level, these relationships are consistent with the hypothesis that an ancestral metabolic-stress system was co-opted during the evolution of worker division of labor. Advancing this framework will require investigating tissue-specific *Crz* signaling, endogenous peptide release, the functional consequences of the Crz–IIS connection, and comparative tests across independently evolved social lineages.

## Acknowledgements

We thank Cahit Ozturk for his assistance with bee husbandry. We thank Krishna Parasawar and Nina Zhang of the Office of Research, Innovation and Impact’s Analytical and Biological Mass Spectrometry Core Facility at the University of Arizona (https://www.cores.arizona.edu/facilities/massspec) for performing mass spectrometry analyses. We thank Meet Zandawala for his helpful comments on the manuscript.

## Data and Code Accessibility Statement

The analytical datasets and R code used for the analyses reported in this study are available on Figshare at https://doi.org/10.6084/m9.figshare.34128333.

