## Supplementary Information for "Corazonin links social role and metabolic-stress responses in honey bee workers"

**Supplemental Information**

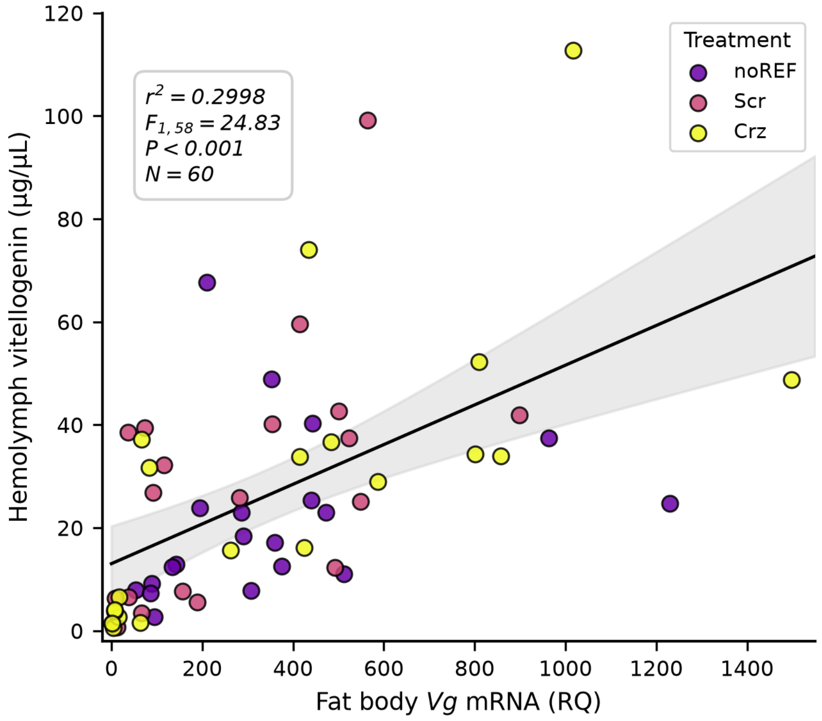


Figure S1. Correlation between relative fat body *Vg* mRNA expression and circulating vitellogenin protein titer. Linear regression of hemolymph vitellogenin protein concentration (µg/µL, measured by SDS-PAGE) on relative fat body *Vg* mRNA abundance (RQ, quantified by RT-qPCR) in individual 7-day-old honey bee workers from the corazonin injection pilot experiment ($N$ = 60; noREF = non-injected reference, Scr = scrambled peptide control, Crz = corazonin peptide injected). Solid line indicates the ordinary least-squares linear regression fit (*titer* = 13.040 + 0.0385 × *Vg*; $r^2$ = 0.2998, $F_{1,58}$ = 24.83, $P$ < 0.001); shaded ribbon denotes the 95% confidence interval.

Table S1. qPCR primer sequences.

| Gene | NCBI Accession # | Forward primer 5\`-3\` | Reverse primer 3\`-5\` | Notes |
|---|---|---|---|---|
| *Actin* | NM_00118514 5.1 | TGCCAACACTGTCCTTTCTG | AGAATTGACCCACCAATCCA | |
| *Crz* | NM_00101296 3.1 | GGAACTTGCGAATAGGAAT G | AAGGCGTTTGAAACAGTTG A | |
| *CrzR* | NM_00114392 | AATGCGTGACCTATGGATTT | ATGAGGATGCCAAGAGGTA | Hits |

| | | | | |
|---|---|---|---|---|
| | 1.1 | | G | dsRNA |
| *CrzR* | NM_001143921.1 | CGAAGATACAATGAAGAACAAC | AGGAATGAAAGGGATGACTC | Redesigned to not hit dsRNA |
| *IRS* | XM_006565041.3 | AGTCATCGTAAGAATAGATTGGA | AGTCATCGTAAGAATAGATTGGA | |
| *UpdlA* | XM_003251746.4 | TTTACCGGAGATTTGGAAGTG | TGTACGTTCTTCTAGCGTAATC | |
| *Cyp314a1* | NM_001040257.1 | AATGGGATTGGAGAGCACAT | GTAACAATTTCCAAAGCGGC | |
| *Vg* | NM_001011578.1 | GTTGGAGAGCAACATGCAGA | TCGATCCATTCCTTGATGGT | |

Table S2. Crz peptide sequences.

| Name | Species | Sequence | Full Sequence |
|---|---|---|---|
| [Thr(4), His(7)]-corazonin | honey bee | pE-TFTYSHGWTN-amide | pGlu-Thr-Phe-Thr-Tyr-Ser-His-Gly-Trp-Thr-Asn-NH2 |
| Randomly_scrambled_91 | honey bee | pEFGNTWHSYTT-amide | pGlu-Phe-Gly-Asn-Thr-Trp-His-Ser-Tyr-Thr-Thr-NH2 |

Table S3. Cloning primers.

| Gene | Cloning direction | Strand | Primer |
|---|---|---|---|
| *CrzR* | Sense | Fwd. | tatagggcgaattgggtaccGTATCTGAGCACGTTCGTGCTTG |
| *CrzR* | Sense | Rev. | tcgaggggggcccggtaccTCATCGTGTAATAGGGCGTCCAC |
| *CrzR* | Antisense | Fwd. | tatagggcgaattgggtaccTCATCGTGTAATAGGGCGTCCAC |

| | | | |
|---|---|---|---|
| *CrzR* | Antisense | Rev. | tcgaggggggcccggtaccGTATCTGAGCACGTTCGTGCTTG |
| *Vg* | Sense | Fwd. | tatagggcgaattgggtaccCGGTCTGACGTCTGACAAATTTG |
| *Vg* | Sense | Rev. | tcgaggggggcccggtaccCCATGCAATGGAACTTGTACGG |
| *Vg* | Antisense | Fwd. | Tatagggcgaattgggtacc CCATGCAATGGAACTTGTACGG |
| *Vg* | Antisense | Rev. | tcgaggggggcccggtaccCGGTCTGACGTCTGACAAATTTG |

**Sense CrzR plasmid sequence:**

ctaaattgtaagcgttaatattttgttaaaattcgcgttaaatttttgttaaatcagctcattttttaaccaataggccgaaatcggcaaaatcc
cttataaatcaaaagaatagaccgagatagggttgagtgttgttccagtttggaacaagagtccactattaaagaacgtggactccaac
gtcaaagggcgaaaaaccgtctatcagggcgatggcccactacgtgaaccatcaccctaatcaagttttttggggtcgaggtgccgta
aagcactaaatcggaaccctaaagggagcccccgatttagagcttgacggggaaagccggcgaacgtggcgagaaaggaagg
gaagaaagcgaaaggagcgggcgctagggcgctggcaagtgtagcggtcacgctgcgcgtaaccaccacacccgccgcgctta
atgcgccgctacagggcgcgtcccattcgccattcaggctgcgcaactgttgggaagggcgatcggtgcgggcctcttcgctattacg
ccagctggcgaaagggggatgtgctgcaaggcgattaagttgggtaacgccagggttttcccagtcacgacgttgtaaaacgacgg
ccagtgagcgcgcgtaatacgactcactatagggcgaattgggtaccCAGCCTGTATCTGAGCACGTTCGTGCTT
GTGCTGATCGGTGTGGACAGGTTCTTTGCCATAAGATATCCTATGAAAGGGATGAACACCGC
TGACAGATGTTTGAAGTTCATCGTCGCGGCTTGGATATTATCCTTTGTACTGTCATTGCCTCA
GATTATCATATTCCGCGTTGTCCAGGGGCCGTTCGTCGAGAAATTCGAGCAATGCGTGACCT
ATGGATTTTACACAGAGCCCTGGCAGGAACAGCTCTACGTATCGTTCGGTTTGTTTTCCATG
TTTCTTCTACCTCTTGGCATCCTCATCGCCACCTACGTTTTCACCATAATTACAATATCTCGGA
GCGAAAGAATGTTCAAAGTGAAATTGGCCAATAACGATATATGCCACGTAAACGGAGATGTA
AATCGAAGGAAATTAATGTATCGAGCAAAAGCTAAATCATTGAGGATCAGTATAGTTATAGTGA
CTGCCTTTATATTTTGGTGGACGCCCTATTACAggtaccgggccccccctcgaggtcgacggtatcgataagctt
gatatcgaattcctgcagcccgggggatccactagttctagagcggccgccaccgcggtggagctccagcttttgttccctttagtgagg
gttaattgcgcgcttggcgtaatcatggtcatagctgtttcctgtgtgaaattgttatccgctcacaattccacacaacatacgagccggaa
gcataaagtgtaaagcctggggtgcctaatgagtgagctaactcacattaattgcgttgcgctcactgcccgctttccagtcgggaaac
ctgtcgtgccagctgcattaatgaatcggccaacgcgcggggagaggcggtttgcgtattgggcgctcttccgcttcctcgctcactgac
tcgctgcgctcggtcgttcggctgcggcgagcggtatcagctcactcaaaggcggtaatacggttatccacagaatcaggggataac
gcaggaaagaacatgtgagcaaaaggccagcaaaaggccaggaaccgtaaaaaggccgcgttgctggcgtttttccataggctcc
gcccccctgacgagcatcacaaaaatcgacgctcaagtcagaggtggcgaaacccgacaggactataaagataccaggcgtttcc
ccctggaagctccctcgtgcgctctcctgttccgaccctgccgcttaccggatacctgtccgcctttctcccttcgggaagcgtggcgcttt
ctcatagctcacgctgtaggtatctcagttcggtgtaggtcgttcgctccaagctgggctgtgtgcacgaaccccccgttcagcccgacc
gctgcgccttatccggtaactatcgtcttgagtccaacccggtaagacacgacttatcgccactggcagcagccactggtaacaggatt
agcagagcgaggtatgtaggcggtgctacagagttcttgaagtggtggcctaactacggctacactagaaggacagtatttggtatctg
cgctctgctgaagccagttaccttcggaaaaagagttggtagctcttgatccggcaaacaaaccaccgctggtagcggtggtttttttgttt
gcaagcagcagattacgcgcagaaaaaaaggatctcaagaagatcctttgatcttttctacggggtctgacgctcagtggaacgaaa
actcacgttaagggattttggtcatgagattatcaaaaaggatcttcacctagatccttttaaattaaaaatgaagttttaaatcaatctaaa
gtatatatgagtaaacttggtctgacagttaccaatgcttaatcagtgaggcacctatctcagcgatctgtctatttcgttcatccatagttgc
ctgactccccgtcgtgtagataactacgatacgggagggcttaccatctggccccagtgctgcaatgataccgcgagacccacgctc
accggctccagatttatcagcaataaaccagccagccggaagggccgagcgcagaagtggtcctgcaactttatccgcctccatcc

agtctattaattgttgccgggaagctagagtaagtagttcgccagttaatagtttgcgcaacgttgttgccattgctacaggcatcgtggtgt
cacgctcgtcgtttggtatggcttcattcagctccggttcccaacgatcaaggcgagttacatgatcccccatgttgtgcaaaaaagcggt
tagctccttcggtcctccgatcgttgtcagaagtaagttggccgcagtgttatcactcatggttatggcagcactgcataattctcttactgtc
atgccatccgtaagatgcttttctgtgactggtgagtactcaaccaagtcattctgagaatagtgtatgcggcgaccgagttgctcttgccc
ggcgtcaatacgggataataccgcgccacatagcagaactttaaaagtgctcatcattggaaaacgttcttcggggcgaaaactctca
aggatcttaccgctgttgagatccagttcgatgtaacccactcgtgcacccaactgatcttcagcatcttttactttcaccagcgtttctgggt
gagcaaaaacaggaaggcaaaatgccgcaaaaaagggaataagggcgacacggaaatgttgaatactcatactcttcctttttcaa
tattattgaagcatttatcagggttattgtctcatgagcggatacatatttgaatgtatttagaaaaataaacaaataggggttccgcgcac
atttccccgaaaagtgccac

**Antisense CrzR plasmid sequence:**

ctaaattgtaagcgttaatattttgttaaaattcgcgttaaatttttgttaaatcagctcattttttaaccaataggccgaaatcggcaaaatcc
cttataaatcaaaagaatagaccgagatagggttgagtgttgttccagtttggaacaagagtccactattaaagaacgtggactccaac
gtcaaagggcgaaaaaccgtctatcagggcgatggcccactacgtgaaccatcaccctaatcaagttttttggggtcgaggtgccgta
aagcactaaatcggaaccctaaagggagcccccgatttagagcttgacggggaaagccggcgaacgtggcgagaaaggaagg
gaagaaagcgaaaggagcgggcgctagggcgctggcaagtgtagcggtcacgctgcgcgtaaccaccacacccgccgcgctta
atgcgccgctacagggcgcgtcccattcgccattcaggctgcgcaactgttgggaagggcgatcggtgcgggcctcttcgctattacg
ccagctggcgaaagggggatgtgctgcaaggcgattaagttgggtaacgccagggttttcccagtcacgacgttgtaaaacgacgg
ccagtgagcgcgcgtaatacgactcactatagggcgaattgggtaccTGTAATAGGGCGTCCACCAAAATATAAAG
GCAGTCACTATAACTATACTGATCCTCAATGATTTAGCTTTTGCTCGATACATTAATTTCCTTC
GATTTACATCTCCGTTTACGTGGCATATATCGTTATTGGCCAATTTCACTTTGAACATTCTTTC
GCTCCGAGATATTGTAATTATGGTGAAAACGTAGGTGGCGATGAGGATGCCAAGAGGTAGAA
GAAACATGGAAAACAAACCGAACGATACGTAGAGCTGTTCCTGCCAGGGCTCTGTGTAAAA
TCCATAGGTCACGCATTGCTCGAATTTCTCGACGAACGGCCCCTGGACAACGCGGAATATG
ATAATCTGAGGCAATGACAGTACAAAGGATAATATCCAAGCCGCGACGATGAACTTCAAACAT
CTGTCAGCGGTGTTCATCCCTTTCATAGGATATCTTATGGCAAAGAACCTGTCCACACCGAT
CAGCACAAGCACGAACGTGCTCAGATACAGGCTGggtaccgggccccccctcgaggtcgacggtatcgata
agcttgatatcgaattcctgcagcccgggggatccactagttctagagcggccgccaccgcggtggagctccagcttttgttccctttagt
gagggttaattgcgcgcttggcgtaatcatggtcatagctgtttcctgtgtgaaattgttatccgctcacaattccacacaacatacgagcc
ggaagcataaagtgtaaagcctggggtgcctaatgagtgagctaactcacattaattgcgttgcgctcactgcccgctttccagtcggg
aaacctgtcgtgccagctgcattaatgaatcggccaacgcgcggggagaggcggtttgcgtattgggcgctcttccgcttcctcgctca
ctgactcgctgcgctcggtcgttcggctgcggcgagcggtatcagctcactcaaaggcggtaatacggttatccacagaatcagggg
ataacgcaggaaagaacatgtgagcaaaaggccagcaaaaggccaggaaccgtaaaaaggccgcgttgctggcgtttttccata

ggctccgccccccctgacgagcatcacaaaaatcgacgctcaagtcagaggtggcgaaacccgacaggactataaagataccagg
cgtttccccctggaagctccctcgtgcgctctcctgttccgaccctgccgcttaccggatacctgtccgcctttctcccttcgggaagcgtg
gcgctttctcatagctcacgctgtaggtatctcagttcggtgtaggtcgttcgctccaagctgggctgtgtgcacgaaccccccgttcagc
ccgaccgctgcgccttatccggtaactatcgtcttgagtccaacccggtaagacacgacttatcgccactggcagcagccactggtaa
caggattagcagagcgaggtatgtaggcggtgctacagagttcttgaagtggtggcctaactacggctacactagaaggacagtattt
ggtatctgcgctctgctgaagccagttaccttcggaaaaagagttggtagctcttgatccggcaaacaaaccaccgctggtagcggtg
gtttttttgtttgcaagcagcagattacgcgcagaaaaaaaggatctcaagaagatcctttgatcttttctacggggtctgacgctcagtgg
aacgaaaactcacgttaagggattttggtcatgagattatcaaaaaggatcttcacctagatccttttaaattaaaaatgaagttttaaatc
aatctaaagtatatatgagtaaacttggtctgacagttaccaatgcttaatcagtgaggcacctatctcagcgatctgtctatttcgttcatcc
atagttgcctgactccccgtcgtgtagataactacgatacgggagggcttaccatctggccccagtgctgcaatgataccgcgagacc
cacgctcaccggctccagatttatcagcaataaaccagccagccggaagggccgagcgcagaagtggtcctgcaactttatccgcc
tccatccagtctattaattgttgccgggaagctagagtaagtagttcgccagttaatagtttgcgcaacgttgttgccattgctacaggcatc
gtggtgtcacgctcgtcgtttggtatggcttcattcagctccggttcccaacgatcaaggcgagttacatgatcccccatgttgtgcaaaa
aagcggttagctccttcggtcctccgatcgttgtcagaagtaagttggccgcagtgttatcactcatggttatggcagcactgcataattct
cttactgtcatgccatccgtaagatgcttttctgtgactggtgagtactcaaccaagtcattctgagaatagtgtatgcggcgaccgagttg
ctcttgcccggcgtcaatacgggataataccgcgccacatagcagaactttaaaagtgctcatcattggaaaacgttcttcggggcga
aaactctcaaggatcttaccgctgttgagatccagttcgatgtaacccactcgtgcacccaactgatcttcagcatcttttactttcaccag
cgtttctgggtgagcaaaaacaggaaggcaaaatgccgcaaaaaagggaataagggcgacacggaaatgttgaatactcatactc
ttcctttttcaatattattgaagcatttatcagggttattgtctcatgagcggatacatatttgaatgtatttagaaaaataaacaaataggggtt
ccgcgcacatttccccgaaaagtgccac

**Sense Vg plasmid sequence:**

ctaaattgtaagcgttaatattttgttaaaattcgcgttaaatttttgttaaatcagctcattttttaaccaataggccgaaatcggcaaaatcc
cttataaatcaaaagaatagaccgagatagggttgagtgttgttccagtttggaacaagagtccactattaaagaacgtggactccaac
gtcaaagggcgaaaaaccgtctatcagggcgatggcccactacgtgaaccatcaccctaatcaagttttttggggtcgaggtgccgta
aagcactaaatcggaaccctaaagggagcccccgatttagagcttgacggggaaagccggcgaacgtggcgagaaaggaagg
gaagaaagcgaaaggagcgggcgctagggcgctggcaagtgtagcggtcacgctgcgcgtaaccaccacacccgccgcgctta
atgcgccgctacagggcgcgtcccattcgccattcaggctgcgcaactgttgggaagggcgatcggtgcgggcctcttcgctattacg
ccagctggcgaaagggggatgtgctgcaaggcgattaagttgggtaacgccagggttttcccagtcacgacgttgtaaaacgacgg
ccagtgagcgcgcgtaatacgactcactatagggcgaattgggtaccCGGTCTGACGTCTGACAAATTTGACGTA
TCCCTGGCTCTGGACGGTGAACGCGTCATGCTCAAGGCGTCTGAAGATTATCGCTACTCCG
TTCGCGGTCTCTGCGGCAATTTCGACCACGACTCGACCAACGACTTCGTGGGACCCAAGA
ACTGCCTGTTCAGAAAGCCCGAACACTTCGTAGCCAGCTACGCTTTGATCAGTAACCAATG

CGAGGGCGACTCGTTGAACGTGGCGAAATCTCTTCAGGATCACGATTGCATCCGACAGGA
GAGGACCCAGCAGAGAAACGTGATCAGCGACAGCGAATCGGGACGATTGGACACCGAGAT
GTCGACCTGGGGTTACCACCATAACGTTAACAAACATTGCACGATCCACAGGACCCAGGTA
AAGGAGACCGACGACAAGATCTGCTTCACCATGCGTCCAGTGGTCTCTTGCGCGTCTGGTT
GCACCGCCGTCGAGACGAAATCGAAGCCGTACAAGTTCCATTGCATGGggtaccgggccccccctc
gaggtcgacggtatcgataagcttgatatcgaattcctgcagcccgggggatccactagttctagagcggccgccaccgcggtggag
ctccagcttttgttccctttagtgagggttaattgcgcgcttggcgtaatcatggtcatagctgtttcctgtgtgaaattgttatccgctcacaatt
ccacacaacatacgagccggaagcataaagtgtaaagcctggggtgcctaatgagtgagctaactcacattaattgcgttgcgctca
ctgcccgctttccagtcgggaaacctgtcgtgccagctgcattaatgaatcggccaacgcgcggggagaggcggtttgcgtattgggc
gctcttccgcttcctcgctcactgactcgctgcgctcggtcgttcggctgcggcgagcggtatcagctcactcaaaggcggtaatacggtt
atccacagaatcaggggataacgcaggaaagaacatgtgagcaaaaggccagcaaaaggccaggaaccgtaaaaaggccgc
gttgctggcgtttttccataggctccgcccccctgacgagcatcacaaaaatcgacgctcaagtcagaggtggcgaaacccgacagg
actataaagataccaggcgtttccccctggaagctccctcgtgcgctctcctgttccgaccctgccgcttaccggatacctgtccgcctttc
tcccttcgggaagcgtggcgctttctcatagctcacgctgtaggtatctcagttcggtgtaggtcgttcgctccaagctgggctgtgtgcac
gaaccccccgttcagcccgaccgctgcgccttatccggtaactatcgtcttgagtccaacccggtaagacacgacttatcgccactgg
cagcagccactggtaacaggattagcagagcgaggtatgtaggcggtgctacagagttcttgaagtggtggcctaactacggctaca
ctagaaggacagtatttggtatctgcgctctgctgaagccagttaccttcggaaaaagagttggtagctcttgatccggcaaacaaacc
accgctggtagcggtggtttttttgtttgcaagcagcagattacgcgcagaaaaaaaggatctcaagaagatcctttgatcttttctacgg
ggtctgacgctcagtggaacgaaaactcacgttaagggattttggtcatgagattatcaaaaaggatcttcacctagatccttttaaatta
aaaatgaagttttaaatcaatctaaagtatatatgagtaaacttggtctgacagttaccaatgcttaatcagtgaggcacctatctcagcg
atctgtctatttcgttcatccatagttgcctgactccccgtcgtgtagataactacgatacgggagggcttaccatctggccccagtgctgc
aatgataccgcgagacccacgctcaccggctccagatttatcagcaataaaccagccagccggaagggccgagcgcagaagtgg
tcctgcaactttatccgcctccatccagtctattaattgttgccgggaagctagagtaagtagttcgccagttaatagtttgcgcaacgttgtt
gccattgctacaggcatcgtggtgtcacgctcgtcgtttggtatggcttcattcagctccggttcccaacgatcaaggcgagttacatgatc
ccccatgttgtgcaaaaaagcggttagctccttcggtcctccgatcgttgtcagaagtaagttggccgcagtgttatcactcatggttatgg
cagcactgcataattctcttactgtcatgccatccgtaagatgcttttctgtgactggtgagtactcaaccaagtcattctgagaatagtgtat
gcggcgaccgagttgctcttgcccggcgtcaatacgggataataccgcgccacatagcagaactttaaaagtgctcatcattggaaa
acgttcttcggggcgaaaactctcaaggatcttaccgctgttgagatccagttcgatgtaacccactcgtgcacccaactgatcttcagc
atcttttactttcaccagcgtttctgggtgagcaaaaacaggaaggcaaaatgccgcaaaaaagggaataagggcgacacggaaat
gttgaatactcatactcttcctttttcaatattattgaagcatttatcagggttattgtctcatgagcggatacatatttgaatgtatttagaaaaa
taaacaaataggggttccgcgcacatttccccgaaaagtgccac

**Antisense Vg plasmid sequence:**

ctaaattgtaagcgttaatattttgttaaaattcgcgttaaatttttgttaaatcagctcattttttaaccaataggccgaaatcggcaaaatcc
cttataaatcaaaagaatagaccgagatagggttgagtgttgttccagtttggaacaagagtccactattaaagaacgtggactccaac
gtcaaagggcgaaaaaccgtctatcagggcgatggcccactacgtgaaccatcaccctaatcaagttttttggggtcgaggtgccgta
aagcactaaatcggaaccctaaagggagcccccgatttagagcttgacggggaaagccggcgaacgtggcgagaaaggaagg
gaagaaagcgaaaggagcgggcgctagggcgctggcaagtgtagcggtcacgctgcgcgtaaccaccacacccgccgcgctta
atgcgccgctacagggcgcgtcccattcgccattcaggctgcgcaactgttgggaagggcgatcggtgcgggcctcttcgctattacg
ccagctggcgaaagggggatgtgctgcaaggcgattaagttgggtaacgccagggttttcccagtcacgacgttgtaaaacgacgg
ccagtgagcgcgcgtaatacgactcactatagggcgaattgggtaccCCATGCAATGGAACTTGTACGGCTTCGA
TTTCGTCTCGACGGCGGTGCAACCAGACGCGCAAGAGACCACTGGACGCATGGTGAAGCA
GATCTTGTCGTCGGTCTCCTTTACCTGGGTCCTGTGGATCGTGCAATGTTTGTTAACGTTAT
GGTGGTAACCCCAGGTCGACATCTCGGTGTCCAATCGTCCCGATTCGCTGTCGCTGATCAC
GTTTCTCTGCTGGGTCCTCTCCTGTCGGATGCAATCGTGATCCTGAAGAGATTTCGCCACG
TTCAACGAGTCGCCCTCGCATTGGTTACTGATCAAAGCGTAGCTGGCTACGAAGTGTTCGG
GCTTTCTGAACAGGCAGTTCTTGGGTCCCACGAAGTCGTTGGTCGAGTCGTGGTCGAAATT
GCCGCAGAGACCGCGAACGGAGTAGCGATAATCTTCAGACGCCTTGAGCATGACGCGTTC
ACCGTCCAGAGCCAGGGATACGTCAAATTTGTCAGACGTCAGACCGggtaccgggccccccctcga
ggtcgacggtatcgataagcttgatatcgaattcctgcagcccgggggatccactagttctagagcggccgccaccgcggtggagctc
cagcttttgttccctttagtgagggttaattgcgcgcttggcgtaatcatggtcatagctgtttcctgtgtgaaattgttatccgctcacaattcc
acacaacatacgagccggaagcataaagtgtaaagcctggggtgcctaatgagtgagctaactcacattaattgcgttgcgctcactg
cccgctttccagtcgggaaacctgtcgtgccagctgcattaatgaatcggccaacgcgcggggagaggcggtttgcgtattgggcgct
cttccgcttcctcgctcactgactcgctgcgctcggtcgttcggctgcggcgagcggtatcagctcactcaaaggcggtaatacggttatc
cacagaatcaggggataacgcaggaaagaacatgtgagcaaaaggccagcaaaaggccaggaaccgtaaaaaggccgcgtt
gctggcgtttttccataggctccgcccccctgacgagcatcacaaaaatcgacgctcaagtcagaggtggcgaaacccgacaggac
tataaagataccaggcgtttccccctggaagctccctcgtgcgctctcctgttccgaccctgccgcttaccggatacctgtccgcctttctc
ccttcgggaagcgtggcgctttctcatagctcacgctgtaggtatctcagttcggtgtaggtcgttcgctccaagctgggctgtgtgcacg
aaccccccgttcagcccgaccgctgcgccttatccggtaactatcgtcttgagtccaacccggtaagacacgacttatcgccactggc
agcagccactggtaacaggattagcagagcgaggtatgtaggcggtgctacagagttcttgaagtggtggcctaactacggctacac
tagaaggacagtatttggtatctgcgctctgctgaagccagttaccttcggaaaaagagttggtagctcttgatccggcaaacaaacca
ccgctggtagcggtggtttttttgtttgcaagcagcagattacgcgcagaaaaaaaggatctcaagaagatcctttgatcttttctacggg
gtctgacgctcagtggaacgaaaactcacgttaagggattttggtcatgagattatcaaaaaggatcttcacctagatccttttaaattaa
aaatgaagttttaaatcaatctaaagtatatatgagtaaacttggtctgacagttaccaatgcttaatcagtgaggcacctatctcagcgat
ctgtctatttcgttcatccatagttgcctgactccccgtcgtgtagataactacgatacgggagggcttaccatctggccccagtgctgcaa
tgataccgcgagacccacgctcaccggctccagatttatcagcaataaaccagccagccggaagggccgagcgcagaagtggtc
ctgcaactttatccgcctccatccagtctattaattgttgccgggaagctagagtaagtagttcgccagttaatagtttgcgcaacgttgttg

ccattgctacaggcatcgtggtgtcacgctcgtcgtttggtatggcttcattcagctccggttcccaacgatcaaggcgagttacatgatcc
cccatgttgtgcaaaaaagcggttagctccttcggtcctccgatcgttgtcagaagtaagttggccgcagtgttatcactcatggttatggc
agcactgcataattctcttactgtcatgccatccgtaagatgcttttctgtgactggtgagtactcaaccaagtcattctgagaatagtgtatg
cggcgaccgagttgctcttgcccggcgtcaatacgggataataccgcgccacatagcagaactttaaaagtgctcatcattggaaaa
cgttcttcggggcgaaaactctcaaggatcttaccgctgttgagatccagttcgatgtaacccactcgtgcacccaactgatcttcagcat
cttttactttcaccagcgtttctgggtgagcaaaaacaggaaggcaaaatgccgcaaaaaagggaataagggcgacacggaaatgt
tgaatactcatactcttcctttttcaatattattgaagcatttatcagggttattgtctcatgagcggatacatatttgaatgtatttagaaaaata
aacaaataggggttccgcgcacatttccccgaaaagtgccac
